## Supplementary Information for "Regulation of coordinated muscular relaxation by a pattern-generating intersegmental circuit"

Hiramoto *et al*.

**Supplementary Figure legends**

**Supplementary Figure 1. Canon neurotransmitter phenotype and activity timing.**

(a) Expression pattern of GCaMP6s driven by *R91C05-Gal4*. (b) Calcium imaging of Canon neurons. Regions of interest (ROIs) used for the analyses shown in Fig. 1b are indicated by squares. (c) Expression pattern of GFP driven by *Canon-spGal4*. (d) Calcium imaging of Canon neurons and aCC MNs. ROIs used for the analyses shown in Fig. 1c, d and Supplementary Fig. 1e are indicated by squares. (e) Plot of the activity timing of A2-A6 aCC MNs and A3-A5 Canon neurons. Normalized time duration from the onset to the peak of calcium signals in each neuron. Activity onset in A2 aCC is set as time 0.0 and peak time in A6 aCC is set as time 1.0. $n$ = 19 waves from two larvae. The error bars represent standard errors of the mean. (f) Canon neurons (green) extend their axons along the anterior commissures. Anti-HRP (magenta) was used to identify anterior (solid arrows) and posterior (dotted arrows) commissures. (g) Presynaptic sites (Brp, red) of Canon neurons (GFP, green) do not overlap with vGluT concentration (vGluT, blue). (h) The cell body of a Canon neuron (GFP, green) is negative for GABA (magenta). (i) Stacked focal planes showing the expression of postsynaptic (DenMark, magenta) and presynaptic (Syt-GFP, green) markers. Arrows indicate dendrites of Canon neurons. A circle indicates the cell body of a Canon neuron. Lines with square heads indicate dendrites and axons of Wave neurons. White dot lines in (b, d) indicate boundaries of VNC. Scale bars in (a-d, f, and i), 50 µm. Scale bars in (g, h), 10 µm.

**Supplementary Figure 2. Wave neurons play no roles in muscular relaxation.**

(a, b) Anterior Wave neurons (arrows) targeted by *R91C05-Gal4* (a) and *Canon-spGal4* (b). (c, d) Normal peristalses in a dissected *MB120B-spGal4* > *UAS-Kir* larva, in which Wave neurons were specifically inactivated. (c) An image of the body-wall muscles visualized by expression of *Mhc*GFP, focusing on ventral muscles in A2-A3 segments. (d) Kymograph generated from the white line in (c) shows smooth muscular movement in both forward (F) and backward (B) peristalses. $n$ = 11 forward and 7 backward waves from one larva. All peristalses examined were normal. Scale bars in (a, b), 50 µm. Scale bars in (c, d), 500 µm.

**Supplementary Figure 3. Pre- and post-synaptic partners of Canon neurons.**

(a, c) Distribution plots of presynaptic (i.e. upstream) (a) and post-synaptic (i.e. downstream) (c) partners of Canon neurons in A2. Transverse axes indicate the number of upstream or downstream neurons of Canon neurons. (b, d) Synaptic occupation by identified upstream (b) and downstream (d) partners of bilateral pairs of Canon neurons in A2 and A3. Neurons that are not shared or group of neurons that have a total of no more than four synapses are classified as “others”.

**Supplementary Figure 4. Characterization of post-synaptic partners of**

**Canon neurons.**

(a) An image of A06c neurons in dorsal view obtained by MCFO. (b) Anterior view of the A06c neuron reconstructed in the region shown in the white box in (a). (c) The cell bodies of A06c neurons express GABA. (d, e) (d) A06c and Canon neurons form sybGRASP-positive contacts. Syb-sp-GFP1-10 and sp-GFP11 were expressed by *R91C05-LexA* and *A06c-Gal4*, respectively. (e) Anterior view of the sybGRASP signals reconstructed in the region shown in the red box in (d). (f) Plots of calcium signals in aCC MNs and GVLIs during fictive backward locomotion. (g-l) EM reconstructions of intersegmental neurons downstream of Canon neurons. Dorsal (g, i, k) and anterior (h, j, l) view are shown. Scale bars in (a-e), 50 µm.

**Supplementary Figure 5. Reconstruction of Canon in each neuromere.**

(a) EM reconstruction of right-side Canon neurons located in segments A2-A5. (b) A table of percentage of postsynapse occupation between Canon neurons. Canon neurons form synapses with each other bidirectionally.

**Supplementary Figure 6. *Canon-spGal4 > TeTxLC* semi-intact preparations generate complete backward peristalses.**

(a) An image of a dissected larva used for the analysis. The dotted white line indicates the location used to make the kymographs shown in (b, c). (b, c) Kymographs showing the propagation of muscle contraction (arrows with dotted lines) during forward (b) and backward (c) peristalses in control (-*TeTxLC*) and *Canon > TeTxLC* (+*TeTxLC*) larvae. Note that all backward peristalses propagate all the way through in the *Canon-spGal4 > TeTxLC* larva as in the control, although the propagation is slower as observed in the *Canon-spGal4 > Kir* larvae. $n$ = 8 backward peristalses in two control larvae and 9 backward peristalses in five experimental larvae. This indicates that propagation of excitatory drive is intact in *Canon-spGal4 > TeTxLC* larvae. Scale bars in (a, b), 500 µm.

**Supplementary Figure 7.**

(a) Co-innervation of the A31k premotor neuron by Ifb-Bwd and Canon neurons. A circuit diagram showing the connections between inhibitory premotor neurons, A31k and A02e, and higher-order Ifb-Bwd and Canon neurons. A02e in segment (n) receives inputs from Ifb-Bwd in segment (n+1) but not from Canon, whereas A31k in segment (n) receives inputs from Ifb-Bwd in segment (n+1) and Canon in segment (n+1) and (n+2). Arrows with filled heads indicate cholinergic synapses while bar heads indicate inhibitory outputs. (b) A scheme showing sequential and long-lasting activation of A31k by Ifb-Bwd and Cannon neurons. (c) A wiring diagram of the inhibitory premotor neurons downstream of Ifb-Bwd and Canon neurons. (d-h) Longer activation of A31k compared to A02e. Reanalysis of data obtained in a previous study[1] (d, e) Plots of the normalized GCaMP6f calcium signal of A02e (d) and A31k (e) during fictive locomotion. (f, g) Normalized GCaMP6f signals of A02e (f) and A31k (g) located in A2-5 neuromeres during fictive backward locomotion. Dotted lines indicate initiation timing of A02 and A31k located in A2, respectively. In (d-g), wave propagation revealed by RGECO1 signals in A02e neurons was used to determine the motor phase (see Methods). (h) Quantification of normalized activity duration of A02e and A31k in the A3 neuromere. Note that activity duration of A31k is significantly longer than A02e. $n$ = 9 waves in five larvae for A02e and 7 waves in three larvae for A31k. $^{**}p = 5.84 \times 10^{-3} < 0.01$, the two-sided Mann-Whitney U test.

**Supplementary Movie 1. Canon neurons wave like activity with backward**

**direction.**

**Supplementary Movie 2. Canon neurons show wave like activity during fictive backward locomotion.**

**Supplementary Movie 3. A larva stops locomotion immediately when Canon neurons are optogenetically activated.**

**Supplementary Movie 4. Slower muscular relaxation during backward peristalsis of a *Canon-spGal4 > UAS-Kir* animal.**

**Supplementary Movie 5. Termination of Canon activity propagation in a *Canon-spGal4 > UAS-TeTxLC* animal.**

Fig. S1

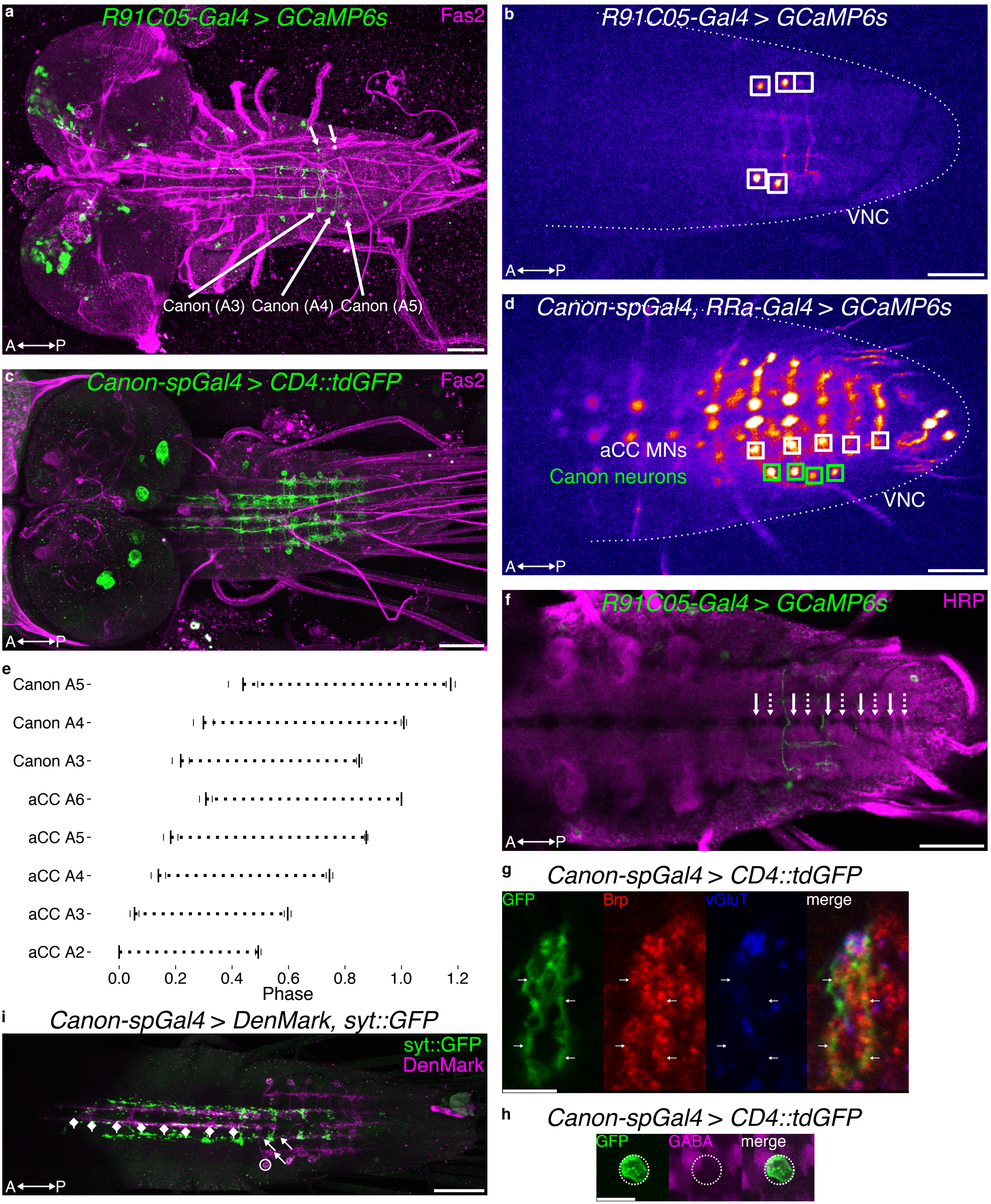

a
R91C05-Gal4 > GCaMP6s
Fas2
Canon (A3)
Canon (A4)
Canon (A5)
A←→P
b
R91C05-Gal4 > GCaMP6s
VNC
A←→P
c
Canon-spGal4 > CD4::tdGFP
Fas2
A←→P
d
Canon-spGal4, RRa-Gal4 > GCaMP6s
aCC MNs
Canon neurons
VNC
A←→P
e
Canon A5
Canon A4
Canon A3
aCC A6
aCC A5
aCC A4
aCC A3
aCC A2
0.0
0.2
0.4
0.6
0.8
1.0
1.2
Phase
f
R91C05-Gal4 > GCaMP6s
HRP
A←→P
g
Canon-spGal4 > CD4::tdGFP
GFP
Brp
vGluT
merge
h
Canon-spGal4 > CD4::tdGFP
GFP
GABA
merge
i
Canon-spGal4 > DenMark, syt::GFP
syt::GFP
DenMark
A←→P

Fig. S2

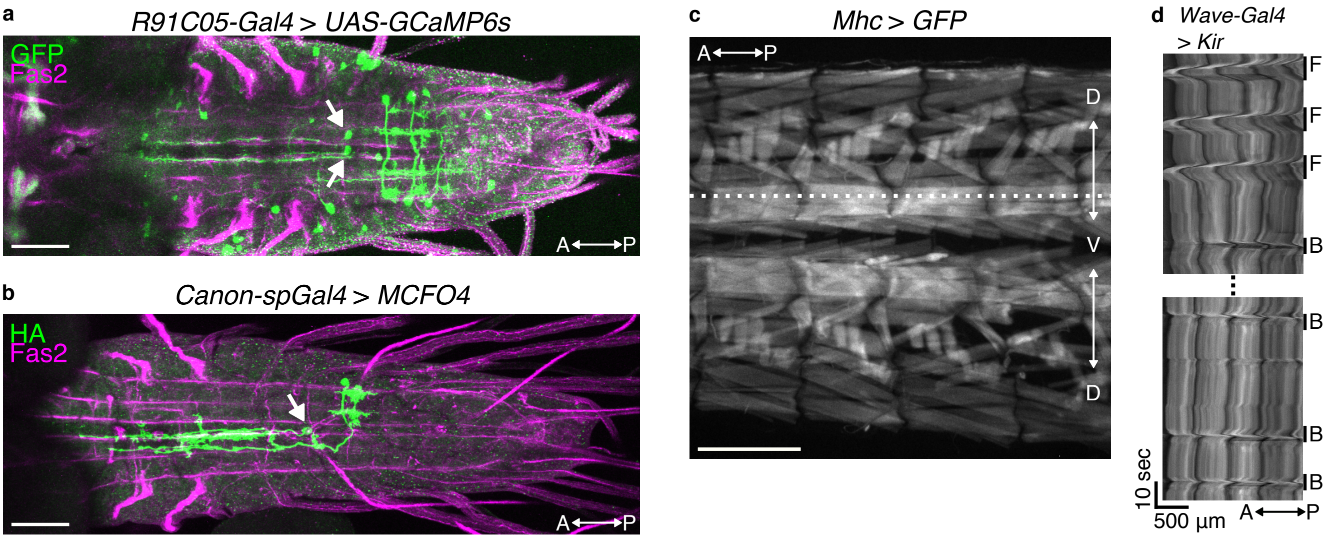
a
R91C05-Gal4 > UAS-GCaMP6s
GFP
Fas2
A P
b
Canon-spGal4 > MCFO4
HA
Fas2
A P
c
Mhc > GFP
A P
D
V
D
d
Wave-Gal4
> Kir
F
F
F
B
B
B
B
10 sec
500 µm
A P

Fig. S3

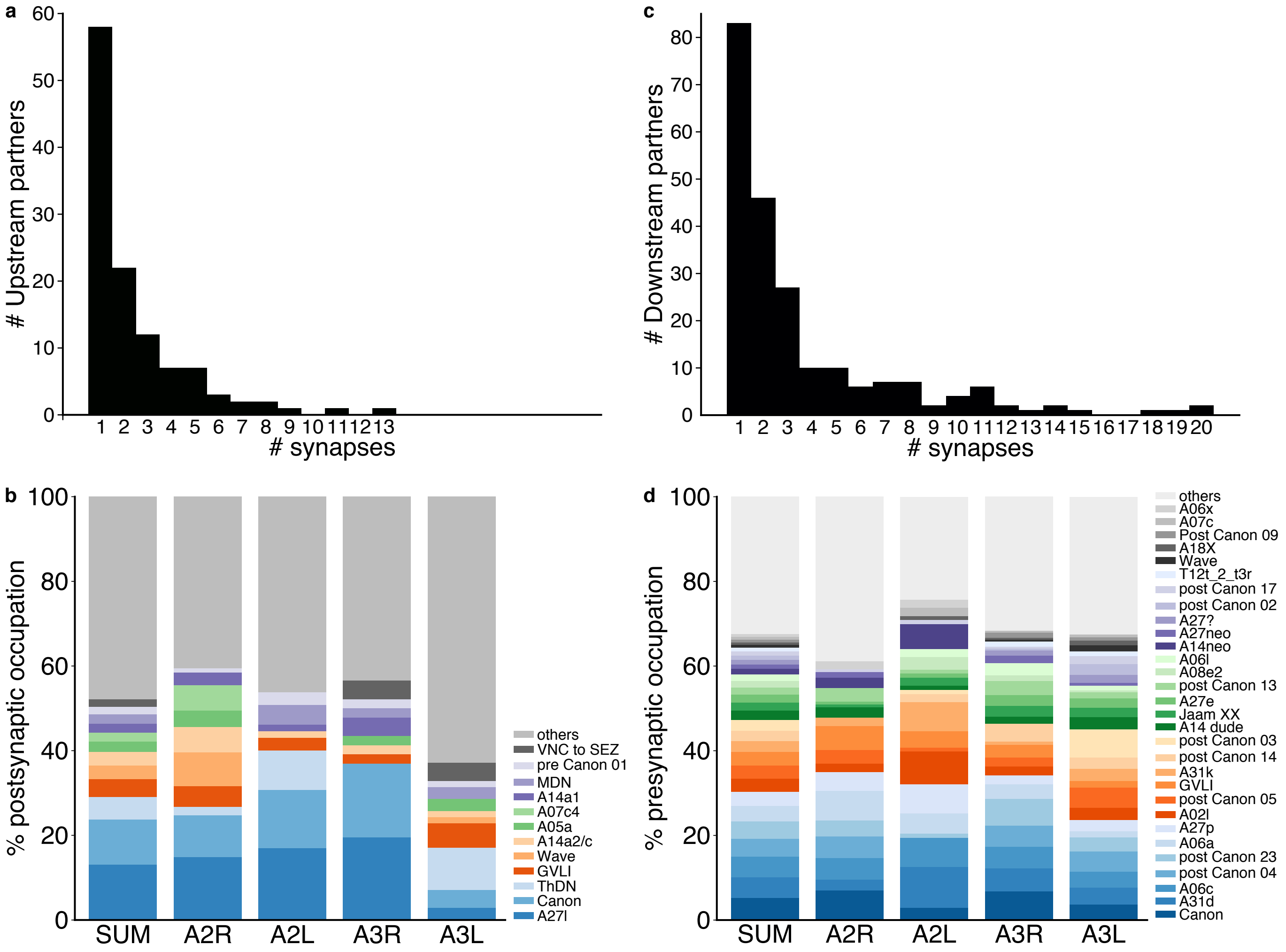
a
### Upstream partners
### synapses
c
### Downstream partners
### synapses
b
% postsynaptic occupation
SUM
A2R
A2L
A3R
A3L
others
VNC to SEZ
pre Canon 01
MDN
A14a1
A07c4
A05a
A14a2/c
Wave
GVLI
ThDN
Canon
A27l
d
% presynaptic occupation
SUM
A2R
A2L
A3R
A3L
others
A06x
A07c
Post Canon 09
A18X
Wave
T12t_2_t3r
post Canon 17
post Canon 02
A27?
A27neo
A14neo
A06l
A08e2
post Canon 13
A27e
Jaam XX
A14 dude
post Canon 03
post Canon 14
A31k
GVLI
post Canon 05
A02l
A27p
A06a
post Canon 23
post Canon 04
A06c
A31d
Canon

Fig. S4

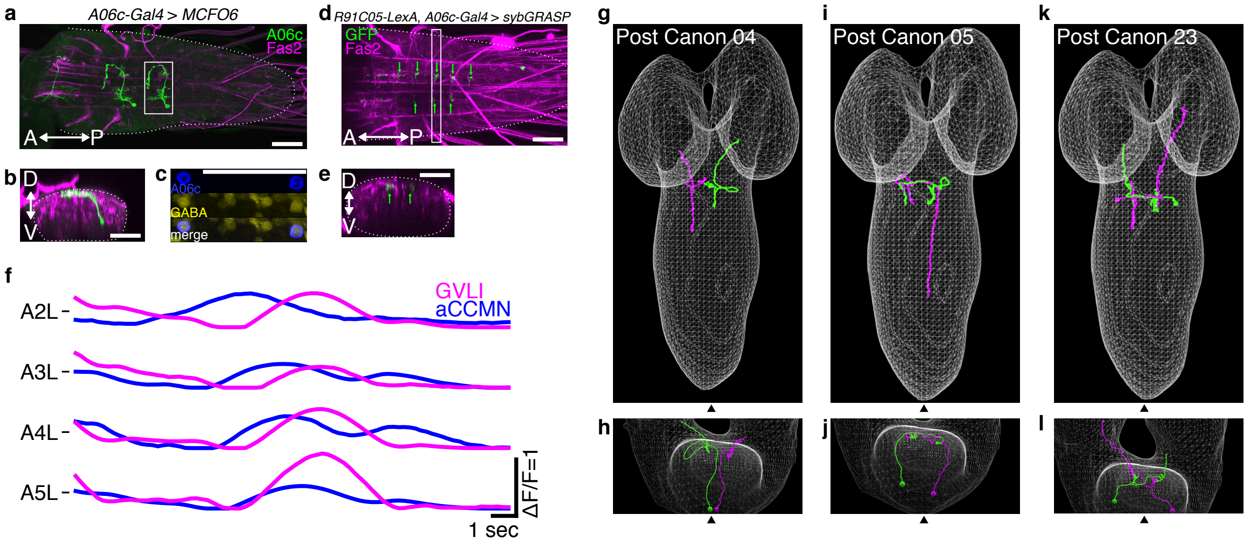

a
A06c-Gal4 > MCFO6
A06c
Fas2
A P
b
D
V
c
A06c
GABA
merge
d
R91C05-LexA, A06c-Gal4 > sybGRASP
GFP
Fas2
A P
e
D
V
f
GVLI
aCCMN
A2L
A3L
A4L
A5L
ΔF/F=1
1 sec
g
Post Canon 04
h
i
Post Canon 05
j
k
Post Canon 23
l

Fig. S5

**a**

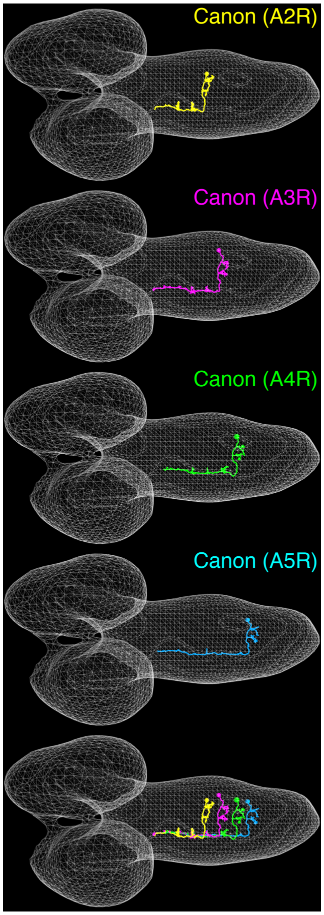


**b**

| Pre \ Post | A2R | A3R | A4R | A5R |
|---|---|---|---|---|
| A2R | 0 | 4.3 | 10 | 2.0 |
| A3R | 4.0 | 0 | 10 | 6.0 |
| A4R | 3.0 | 11 | 0 | 4.0 |
| A5R | 1.0 | 2.2 | 7.7 | 0 |

Fig. S6

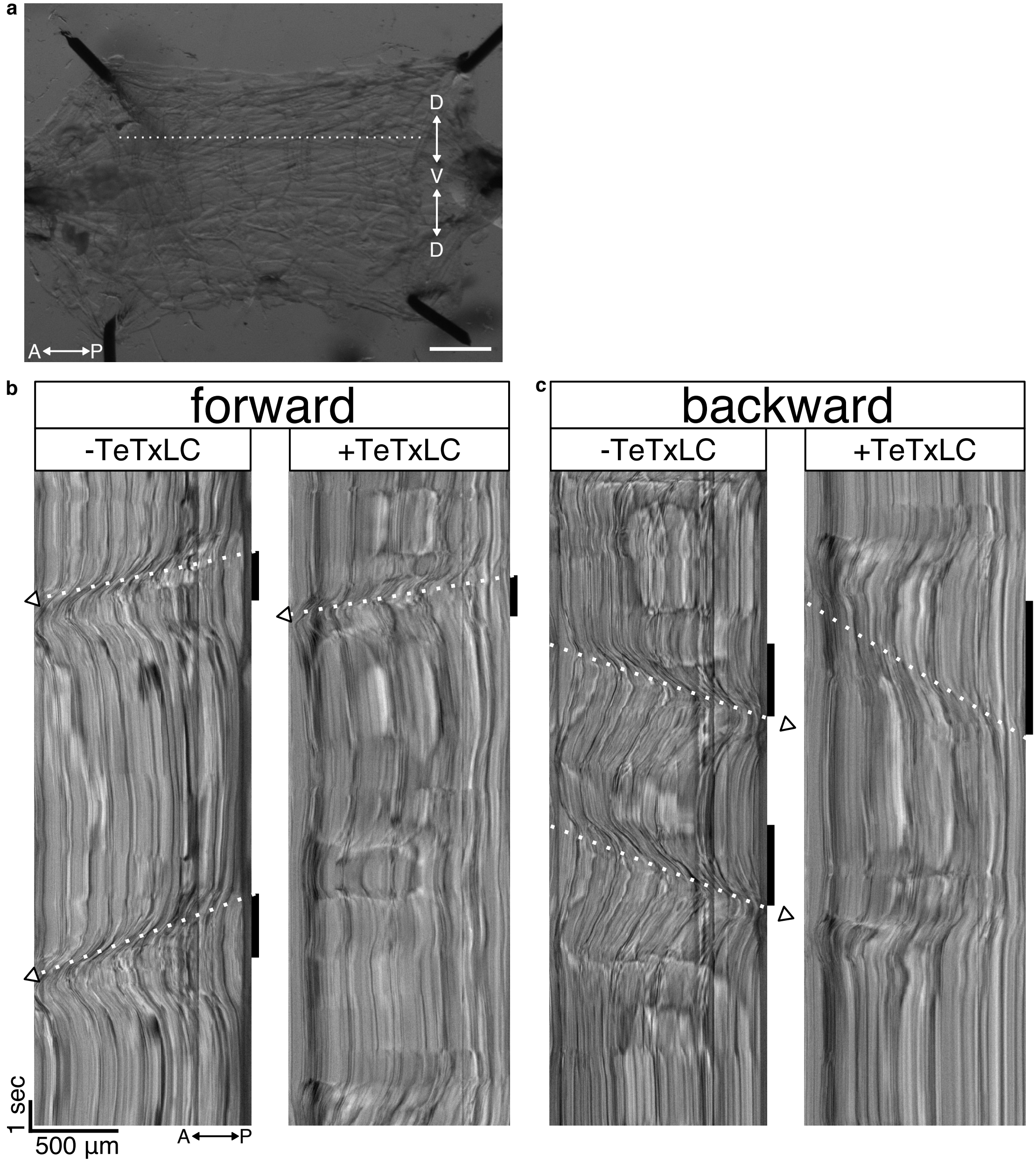
a
D
V
D
A
P
b
forward
-TeTxLC
+TeTxLC
c
backward
-TeTxLC
+TeTxLC
1 sec
500 µm
A
P

Fig. S7

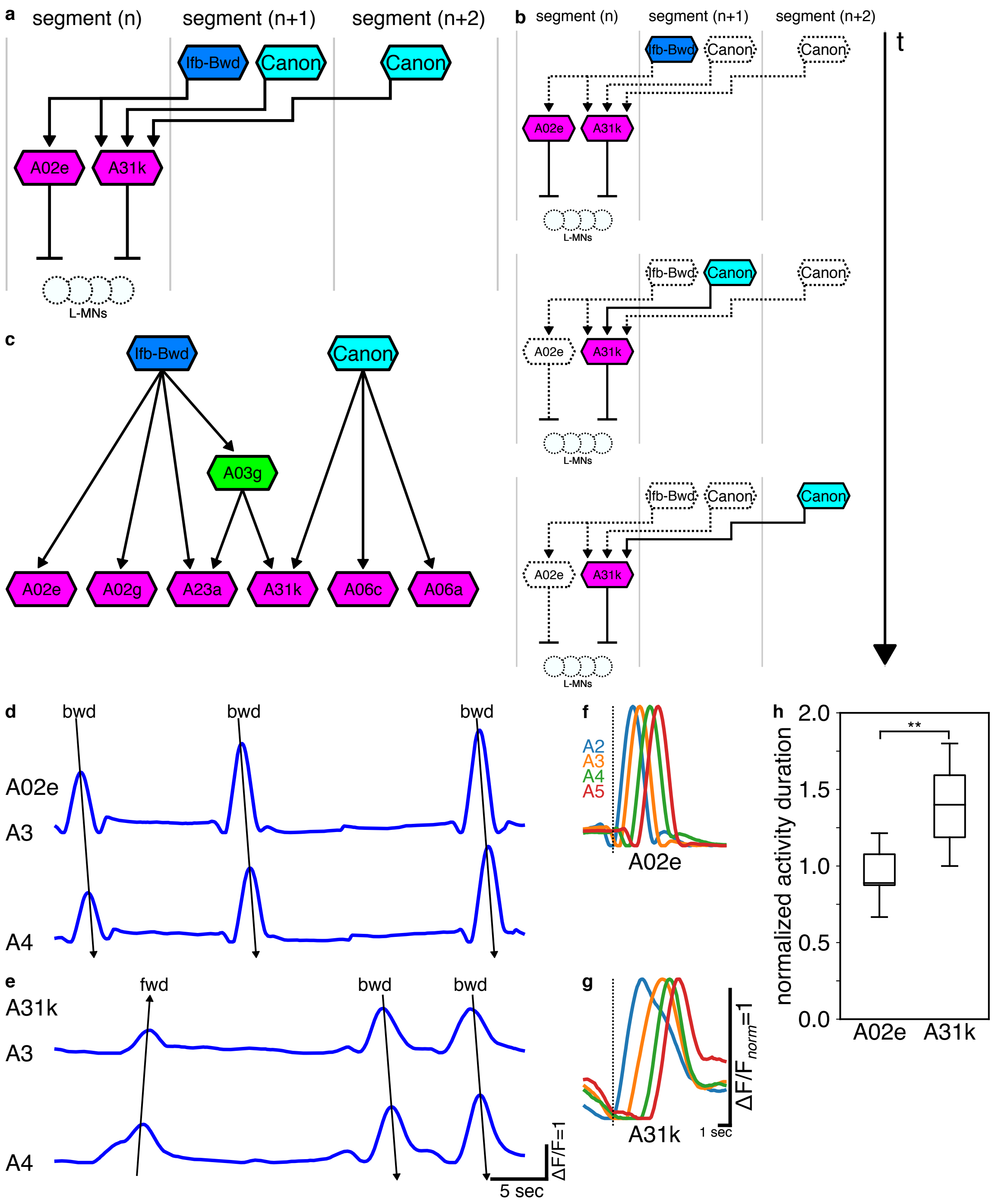

a
segment (n)
segment (n+1)
segment (n+2)
lfb-Bwd
Canon
Canon
A02e
A31k
L-MNs
b
segment (n)
segment (n+1)
segment (n+2)
t
c
lfb-Bwd
Canon
A03g
A02e
A02g
A23a
A31k
A06c
A06a
d
bwd
bwd
bwd
A02e
A3
A4
e
fwd
bwd
bwd
A31k
A3
A4
ΔF/F=1
5 sec
f
A2
A3
A4
A5
A02e
g
A31k
1 sec
ΔF/F_norm=1
h
normalized activity duration
2.0
1.5
1.0
0.5
0.0
**
A02e
A31k
